# PyF2F: A robust and simplified fluorophore-to-fluorophore distance measurement tool for Protein interactions from Imaging Complexes after Translocation experiments

**DOI:** 10.1101/2024.02.16.580512

**Authors:** Altair C. Hernandez, Sebastian Ortiz, Laura I. Betancur, Radovan Dojčilović, Andrea Picco, Marko Kaksonen, Baldo Oliva, Oriol Gallego

## Abstract

**Summary:** Structural knowledge of protein assemblies in their physiological environment is paramount to understand cellular functions at the molecular level. Protein interactions from Imaging Complexes after Translocation (PICT) is a live-cell imaging technique for the structural characterization of macromolecular assemblies in living cells. PICT relies on the measurement of the separation between labelled molecules using fluorescence microscopy and cell engineering. Unfortunately, the required computational tools to extract molecular distances involve a variety of sophisticated software that challenge reproducibility and limit their implementation by highly specialised researchers. Here we introduce PyF2F, a Python-based software that provides a workflow for measuring molecular distances from PICT data, with minimal user programming expertise. We used a published dataset to validate PyF2F performance.

## 1 Introduction

Unravelling the structure of macromolecular complexes is necessary to understand molecular functions, interactions and dynamics that explain the mechanisms controlling the cell’s biology. The structure of molecular assemblies can be solved at high resolution by a number of techniques such as x-ray crystallography, NMR, and cryo-electron microscopy. However, these methods are limited to solve only molecular assemblies that have been previously isolated from their physiological environment. This technical requirement prevents capturing functional conformations and structural dynamics that are central to the mechanism of cellular processes.

Fluorescence microscopy offers the unique opportunity to investigate the spatial organisation of biological macromolecules within their functional environment: the cell. Although light diffraction limits the resolution of fluorescence microscopy to 200-300 nm, localization microscopy is a technique that overcomes this restraint by estimating the centroid position of diffraction limited fluorescence spots (**1-3**). With this method, protein-complex subunits can be specifically labelled with fluorescent markers whose position can be determined at high precision (in the range of 20-30 nanometers) (**2**). Subsequently, the separation between two different subunits labelled with distinguishable fluorophores can be estimated from their respective localization. Repetitive and reproducible measurement of the distance between two fluorophores allows for estimating their true separation with up to one nanometre precision (**1,4-10**). Thus, the measure of fluorophore-to-fluorophore distances can provide outstanding information about conformational changes and molecular interactions (**6,8-10**).

In practice, technical constraints limit the implementation of localization microscopy to interrogate protein complex structures *in situ*. Firstly, not all proteins can be observed as distinguishable diffraction-limited spots when labelled with a fluorophore in living cells. Secondly, the intrinsic dynamic nature of cellular processes prevents repetitive and reproducible imaging of the diffraction limited spots. Thirdly, accurate measurements of distances between fluorophores requires the combination of sophisticated pre-processing (i.e., subtraction of background noise, image registration, cell segmentation) and analysis (i.e., feature detection, gaussian fitting, rejection of outliers) of the raw images.

Our group developed PICT (Protein interactions from Imaging Complexes after Translocation) (**11,12**), a live-cell imaging technique that enables 1) the inducible distribution of fluorescently labelled protein complexes in diffraction-limited spots and 2) the repetitive and reproducible imaging to estimate distances with high precision. PICT employs yeast plasma membrane-associated anchoring platforms and the rapamycin-induced heterodimerization of FK506-binding protein (FKBP) and FKBP-rapamycin binding (FRB) domains to recruit the protein complex of interest (tagged with FRB) to the anchoring platform (tagged with FKBP). The anchor and one subunit of the protein complex are labelled with distinguishable fluorophores (i.e. RFP and GFP). Once the recruitment of the protein complex has succeeded, the centroid position in the equatorial plane of the cell for the anchor and the labelled subunit are determined and their separation is estimated with 2 to 5 nm of precision (**12**). The orientation in which the assembly is anchored can be controlled by fusing the FRB tag to different subunits. Thus, PICT allows measuring the distance between the labelled subunit and the anchor when the complex is recruited in different orientations. Integration of the measured distances allows the trilateration of the fluorophores that label the complex subunits using the anchoring platform as a spatial reference point. This approach has been used to reconstruct the architecture of the exocyst and the conserved oligomeric Golgi (COG) complexes in living cells (**12**).

Unfortunately, the image pre-processing and analysis workflows necessary for a precise distance estimation in PICT still require the installation and version compatibility control of multiple software such as FIJI/ImageJ (**13**), MATLAB (**14**) and R (**15**). In practice, the broad implementation of the PICT method to resolve the architecture of protein assemblies suffers from optimal integration of multiple software and custom scripts and the requirement of specialised expertise in microscopy and image analysis that is rarely found in structural biology laboratories.

Here, we present PyF2F (Python-based Fluorophore-to-Fluorophore ruler), a Python-based open-source software for measuring distances between two fluorescent markers from diffraction-limited microscopy images obtained by PICT. We improved the performance of yeast cell segmentation by taking advantage of pre-trained convolutional neural network (CNN) weights used in YeastSpotter (**16**). PyF2F is an accessible and robust tool that facilitates reproducibility of fluorophore-to-fluorophore distance measurement in living cells. Together with the code and documentation, we provide a walkthrough tutorial that can be easily followed by users unfamiliar with this image analysis protocol. PyF2F can run standalone with a local installation or online using a Colab notebook (https://colab.research.google.com/drive/1kSOnZdwRb4xuznyQIpRNWUBBFKms91M8?usp=sharing).

## MATERIAL & METHODS

### Software architecture

PyF2F has been implemented in Python 3.7 and runs either standalone in any Linux or MacOS terminal or in a Google Colab notebook. The standalone version has dependencies on the following packages:

- *sci-kit image v. 0*.*19*.*2* (**17**)
- *pymicro v*. 0.5.1 (https://www.github.com/heprom/pymicro)
- *vtk v. 9*.*1*.*0* (**18**)
- *trackpy* v. 0.5.0 (**19**)
- *scipy v. 1*.*7*.*3* (**20**)
- *numpy v. 1*.*21*.*2* (**21**)
- *pandas v. 1*.*3*.*5* (**22**)
- *plotly v. 5*.*3*.*1* (https://plot.ly)
- *seaborn v. 0*.*12*.*2* (**23**)
- *matplotlib* v. *3*.*5*.*1* (**24**)
- *h5py v. 2*.*10*.*0 (***25**)
- *keras v. 2*.*1*.*6 (***26**)
- *lmfit v. 1*.*0*.*3 (***27**)
- *opencv_python v*.*4*.*5*.*5*.*62 (***28**)
- *pillow v*.*9*.*5*.*0 (***29**)
- *tensorflow v*.*1*.*15*.*0 (***30**)

PyF2F also integrates customised Python functions of the YeastSpotter for an accurate yeast cell segmentation (**16**).

### Workflow

PyF2F’s workflow consists of four major steps: 1) image registration, 2) image pre-processing and spot-pair detection, 3) spot-pair filtering and 4) distance estimation. We provide recommended parameters given a system equipped with a 100x 1.49 NA objective lens and a camera with a pixel size of 6.45 μm.

#### 1. Image Registration

Before embarking on the measurement of the separation between differentially labelled biomolecules, PyF2F measures and corrects the chromatic aberration intrinsic to the employed microscope setup. As a standard, PyF2F uses one dataset of two-channel images of TetraSpeck beads to correct the chromatic aberration (**6**). PyF2F utilises *pymicro*’s point set registration function (*compute_affine_transform*) (https://github.com/heprom/pymicro) to calculate the registration map and correct the beads’ coordinates for the chromatic aberration. The registration map is computed by applying an affine transformation, which corrects the beads’ coordinates of channel 1 (red-fluorescence signal, which we represent as magenta to facilitate visualisation) using the beads’ coordinates of channel 2 (green-fluorescence signal) as reference (Fig 1. A).

**Fig. 1.**
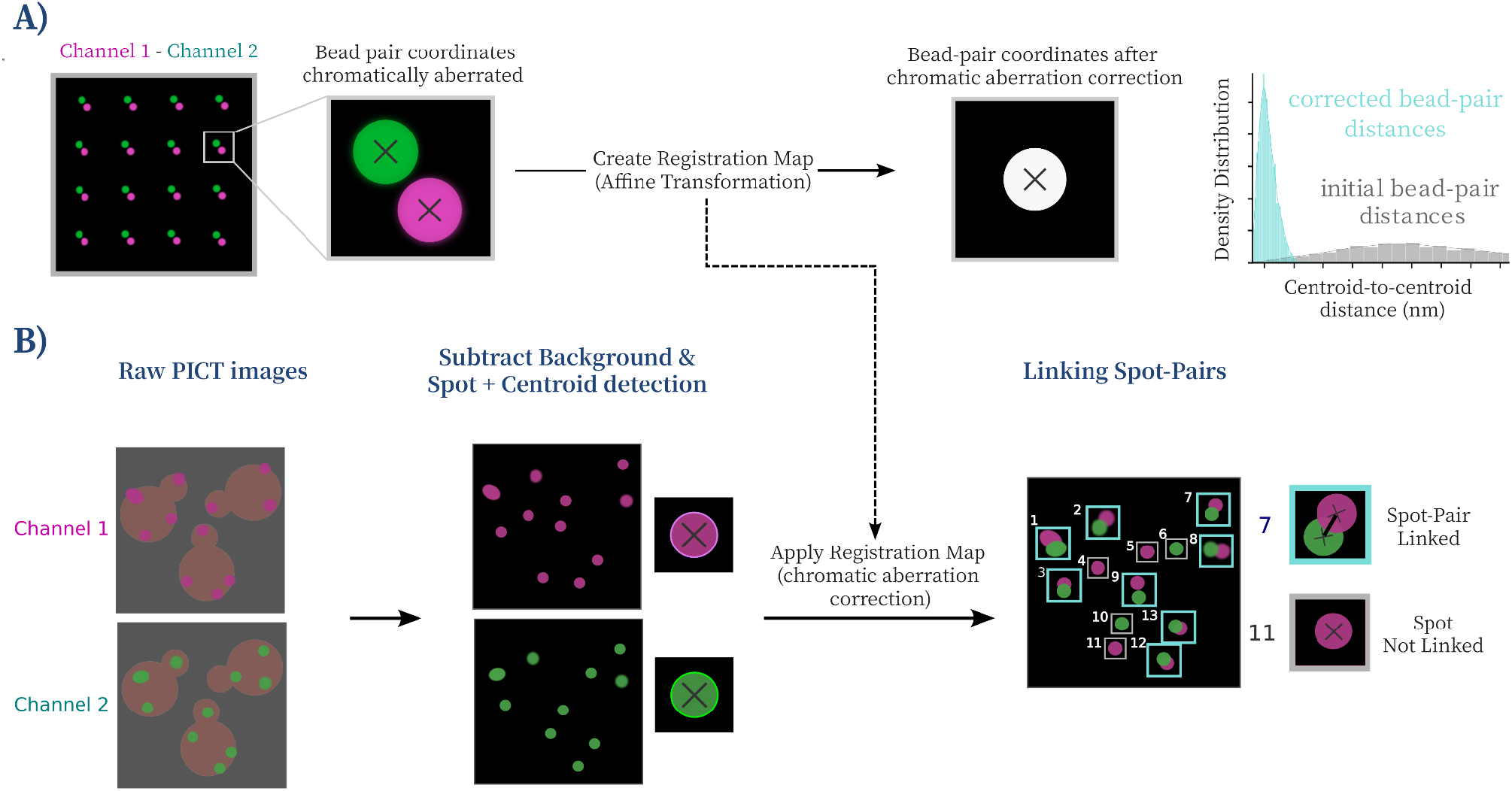
Scheme of Image Registration, Image Pre-processing and Spot Detection. (**A**) **Image registration:** a dataset of two-channel images of TetraSpeck beads (red fluorescence, Ch1-magenta; green fluorescence, Ch2-green) (left). The zoom-in shows the centroid coordinates of representative fluorescent beads for each channel before (centre) and after (right) being corrected for the chromatic aberration using an affine transformation. This allows creating the registration map. As control, the centroid-to-centroid Euclidean distances between two-channel bead pairs are measured before (grey) and after (turquoise) the chromatic aberration correction (right). (**B**) **Image pre-processing:** two-channel images obtained in a PICT experiment are firstly pre-processed to remove the extracellular and cytoplasmic background fluorescence signal (left). **Spot-pair detection**: the centroid coordinates of the fluorescent spot-pairs corresponding to the labelled protein (green) and anchor site (magenta) are localised (zoom-in - centre), linked and corrected for chromatic aberration using the registration map computed with the bead images (right).

#### 2. Image Pre-processing and Spot-Pair Detection

After image registration, the background fluorescence of the images is corrected to localise the centroid of emission of fluorescent markers. Firstly, the extracellular background intensity is subtracted using the *rolling ball* algorithm of the sci-kit image package with a rolling ball radius slightly larger than the cell size. Secondly, to correct the uneven cytoplasmic fluorescence signal at the edge of the cell, PyF2F subtracts the median filtered image from the previous background-subtracted image. The median filtered image is computed using a radius equal to twice the diffraction limit, which is sufficiently large to consistently erase the fluorescent spots from the image while preserving the cell contour (**31**) (Fig. 1. B). For the microscopy setup used as example, we recommend using a rolling ball radius equal to 70 pixels and a median radius of 11 pixels.

Subsequently, PyF2F performs the spot detection on the two channels independently using *trackpy* (**18**). Detected spots (red fluorophore Ch1-magenta, green fluorophore Ch2-green) are then linked within a maximum separation distance between centroid coordinates set by the user. For the microscopy setup used as example, we recommend using a maximum separation distance between 2 to 3 pixels for intra-assembly and inter-assembly distance measurements, respectively. Only linked spot-pairs are selected for posterior analysis. Finally, the coordinates of selected spots in channel 1 are corrected for the chromatic aberration using the registration map saved in step 1 (Fig 1. B).

#### 3. Spot-Pair Filtering

At this step, PyF2F applies a sequence of filters to reject noisy centroid positions of detected spots to ensure that only reliable spot-pairs are selected. Detected spot-pairs are sorted according to the following criteria:

- Isolated events: PyF2F filters out overlapping spot-pairs based on a maximum closest-neighbour distance cut-off. We recommend using a maximum closest-neighbour distance equal to twice the diffraction limit minus one pixel (i.e. 10 pixels for the microscopy setup used as example).
- Distance to the cell contour: as PICT relies on the employment of plasma membrane-associated anchoring platforms, detected spot-pairs must be located at the edge of the cell’s equatorial plane (cell contour). PyF2F utilises a neural-network-based cell segmentation algorithm (**16**) to draw the cell contour, and it selects only those spot coordinates that fall close to the cell contour based on a distance cut-off (Fig 2. A). We recommend using a distance-to-cell-contour cut-off equivalent to twice the diffraction limit plus two pixels (i.e. 13 pixels for the microscopy setup used as example).
- High-quality Spot-pairs: filtering of high quality spot-pairs is done based on the highest dense group of spot-pairs sharing similar properties in brightness (second momentum) and roundness (eccentricity). (Fig 2. A). Spot-pairs with higher probability to be found are selected. We recommend using a cut-off of 0.5 to select the spot-pairs that have a probability of 50% or more to be found (i.e. the 50% denser spot-pair population).
- In-focus Spot-pairs: the distribution of the pixel intensity in each spot is evaluated with a 2D Gaussian fitting. PyF2F evaluates the quality of the interpolation (*R*^*2*^) and selects spot-pairs above the threshold set by the user (0 - 1) (Fig 2. A). We recommend using a threshold of 0.35 to reject spot-pairs with an inappropriate 2D-Gaussian intensity profile.

**Fig. 2.**
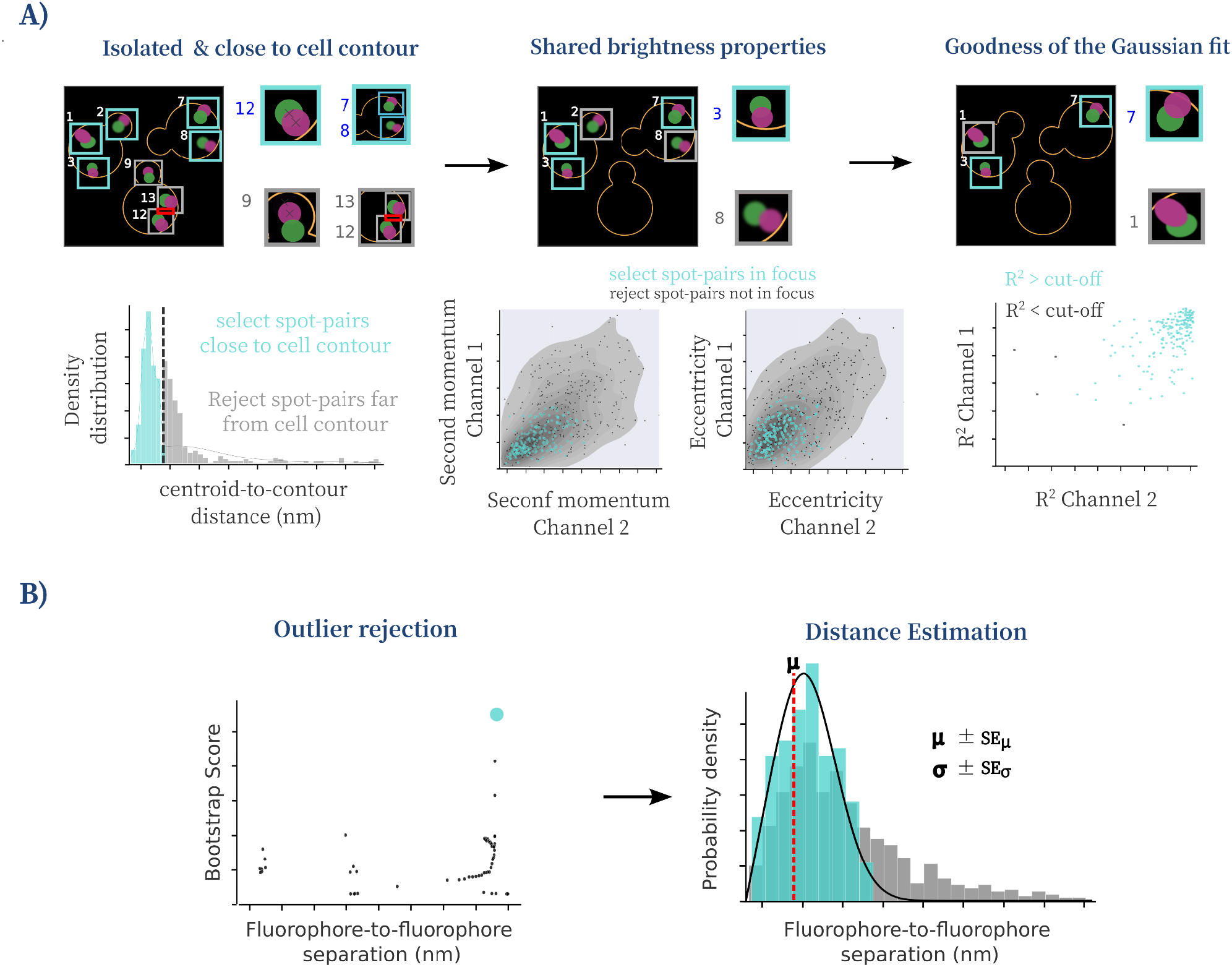
Scheme of Spot-pair Selection, Outlier Rejection and Distance Estimation. (**A**) **Spot-pair Selection**: the contour of the cell is approximated using the CNN weights of YeastSpotter (**16**). Detected spot-pairs are sorted according to their distance to the closest neighbour spot and distance-to-cell contour (left). Only spot-pairs with similar brightness (second momentum) and roundness (eccentricity) are selected (centre) and fitted to a 2D Gaussian distribution (right). The zoom-in illustrates each selection step. Turquoise and grey squares show selected and rejected spot-pairs, respectively. (**B**) **Outlier Rejection and Distance Estimation:** outliers are rejected using a bootstrap method (left, see Supplementary Note S1) to determine the most scored distance distribution without outliers (turquoise dot) that maximises the likelihood for the estimate of μ and σ. The distance distribution is modelled by a Rician distribution (**5**) with a Maximum Likelihood estimate (MLE) to approximate the true separation between fluorophores (μ - red dashed line) and the variance (σ) of the distribution (right). SEμ and Seσ are the respective standard errors of μ and σ estimations.

#### 4. Distance Estimation

The last step estimates the true separation between the two fluorophore labels. The distribution of measured distances *d* that separates paired fluorescent spots follows a Rician distribution (**4,5**) defined as:

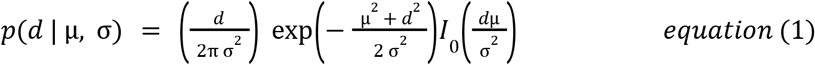

where μ is the true separation between the fluorophores (the one we want to estimate), σ is the distribution variance, and *I*_0_ is the modified Bessel function of integer order 0. The true separation μ can thus be computed with a Maximum Likelihood Estimate (MLE) (**5**). Due to the skewed nature of the distribution, outliers, especially if in the tail of the distribution, can fail the MLE. To reject these outlier measures we implemented a bootstrap method (Fig 2. B, see Supplementary Note S1).

## RESULTS

To evaluate the PyF2F performance, we used a PICT dataset where the hetero-octameric exocyst complex had been recruited to anchoring platforms labelled with RFP (Supplementary Table S1) (**12**). The dataset included images of cells where the exocyst had been anchored in 8 different orientations (i.e. the FRB tag had been fused to a different exocyst subunit). Including all the orientations, we compared a total of 78 intra-assembly distance estimations: 1) 45 distances between the anchor and the 3xGFP tag fused to the C-terminus of exocyst subunits (Fig 3. A), 2) 33 distances between the anchor and the GFP tag fused to the N-terminus of exocyst subunits (Fig 3. B). We also compared 6 inter-assembly distance estimations (distances between the anchor and the GFP fused to Sec2 C-terminus, a protein on the surface of the secretory vesicle that is bound by the exocyst) (Fig 3. C, Supplementary Table S1). A linear fitting showed a strong correlation between the distance estimations (R-squared = 0.93, p-value = 1.1x10^-6^), and the estimated slope (0.97 ± 0.03) and the intercept (-0.26 ± 0.03) suggest that the results obtained with PyF2F are coherent and reproducible with respect to the published dataset (Fig 3. D).

**Fig. 3.**
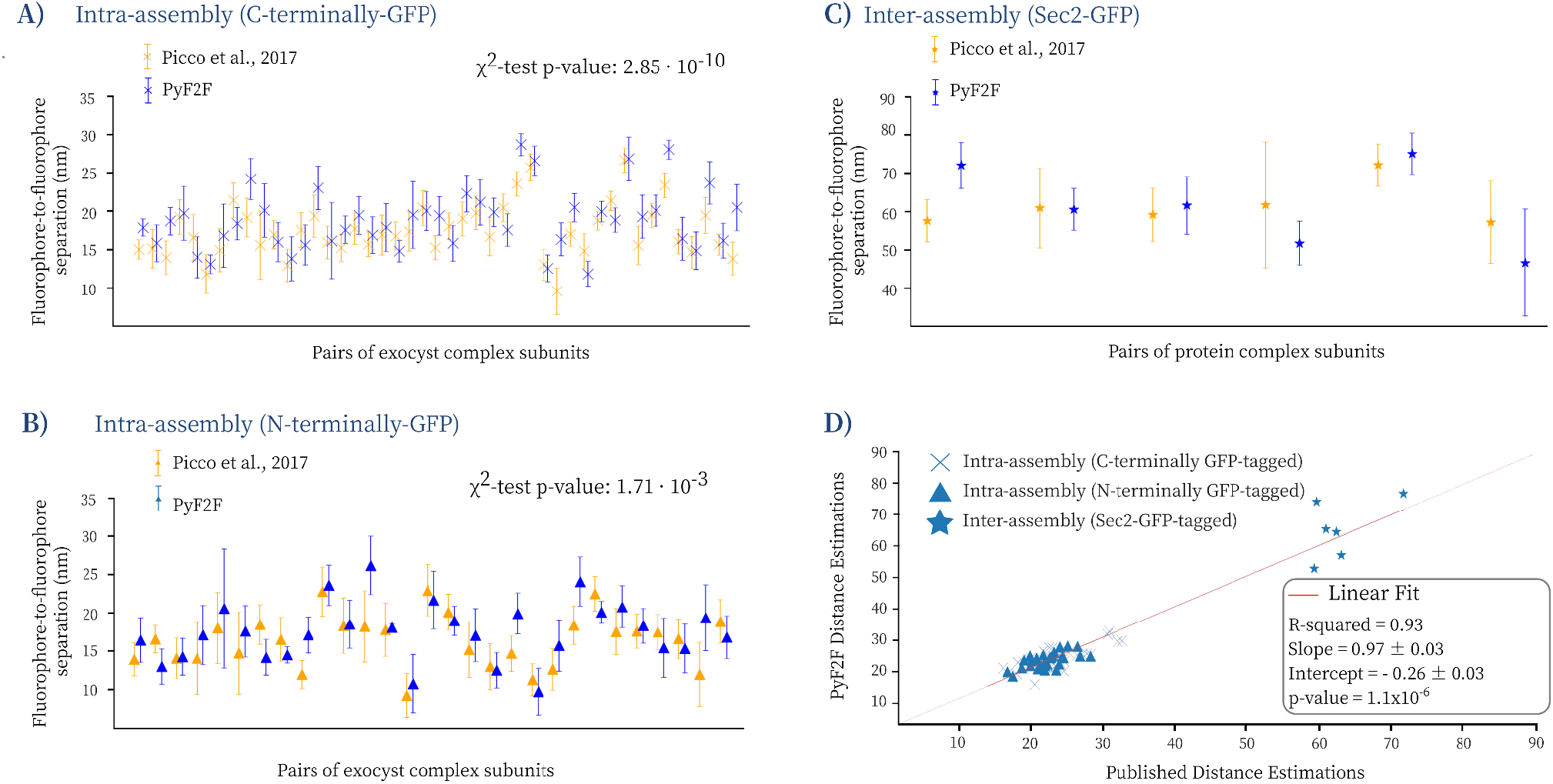
PyF2F validation. Validation of the PyF2F performance by comparing published distance estimations (orange, **12**) with the estimations obtained with PyF2F (blue) (see Supplementary Table S1). (**A**) 45 distances between the anchor and exocyst subunits tagged to 3xGFP at their C-terminus, (**B**) 33 distances between the anchor and exocyst subunits tagged to GFP at their N-terminus and (**C**) 6 distances between the anchor and the GFP fused to Sec2 C-terminus (inter-assembly distances). The difference between the groups of distance estimations was evaluated with a Chi-squared test. (**D**) The correlation between the published and the PyF2F distance estimations was evaluated with a linear fitting (R-squared = 0.93; Slope = 0.97 ± 0.03; Intercept = -0.26 ± 0.03; p-value = 1.1x10-6, n = 84 distances).

We also used a Chi-squared test to corroborate the reproducibility of the measurements obtained by PyF2F. We excluded the Sec2-GFP dataset since the number of measurements was not large enough to perform a reliable Chi-square test. The Chi-squared test showed no significant differences between the intra-assembly distance estimations (p-value of 2.85x10^-10^ and 1.7x10^-3^ for the C-terminal and N-terminal intra-assembly distance measurements respectively) (see Fig. 3. A-B). Overall, the coefficient of determination R^2^ of the linear regression and the Chi-square test indicate that PyF2F delivers reliable distance estimations from PICT measurements.

## DISCUSSION

PyF2F is a Python-based software that requires minimal user programming expertise for estimating the separation between two fluorescent markers imaged with diffraction-limited microscopy in living cells. PyF2F overcomes the limitations of existing approaches by providing a unified application that allows gathering spatial information of proteins translocated to engineered anchoring platforms (with a precision of 2 to 5 nm). Compared to former software (**12**), PyF2F is robust, portable and includes improvements in the yeast cell segmentation (**16**). We provide the required tutorials to facilitate the use of PyF2F by other laboratories (https://colab.research.google.com/drive/1kSOnZdwRb4xuznyQIpRNWUBBFKms91M8?usp=sharing). In combination with the PICT technique, PyF2F generalises the *in situ* structural measurements within and between protein complexes.

## Data Availability

The code and data underlying this article are available on GitHub at https://github.com/GallegoLab/PyF2F under the MIT licence.

## Funding

This work was supported by the Spanish funding agency [PID2021-127773NB-I00/FEDER UE., PRE 2019-088514]; the Unidad de Excelencia Maria de Maeztu [MDM-2014-0370., MCIN/AEI/10.13039/501100011033]; and the Human Frontiers Science Program [RGP0017/2020].

## Conflict of Interest

none declared.

## Notes

https://github.com/GallegoLab/PyF2F

https://zenodo.org/records/8014795

### Competing Interest Statement

The authors have declared no competing interest.

## REFERENCES

1. Mortensen, K. I., Sung, J., Flyvbjerg, H., & Spudich, J. A. (2015). Optimized measurements of separations and angles between intra-molecular fluorescent markers. Nature communications, 6(1), 8621.

2. Thompson, R. E., Larson, D. R., & Webb, W. W. (2002). Precise nanometer localization analysis for individual fluorescent probes. Biophysical journal, 82(5), 2775–2783.

3. Yildiz, A., & Selvin, P. R. (2005). Fluorescence imaging with one nanometer accuracy: application to molecular motors. Accounts of chemical research, 38(7), 574–582.

4. Churchman, L. S., Ökten, Z., Rock, R. S., Dawson, J. F., & Spudich, J. A. (2005). Single molecule high-resolution colocalization of Cy3 and Cy5 attached to macromolecules measures intramolecular distances through time. Proceedings of the National Academy of Sciences, 102(5), 1419–1423.

5. Churchman, L. S., Flyvbjerg, H., & Spudich, J. A. (2006). A non-Gaussian distribution quantifies distances measured with fluorescence localization techniques. Biophysical journal, 90(2), 668–671.

6. Niekamp, S., Sung, J., Huynh, W., Bhabha, G., Vale, R. D., & Stuurman, N. (2019). Nanometer-accuracy distance measurements between fluorophores at the single-molecule level. Proceedings of the National Academy of Sciences, 116(10), 4275–4284.

7. Pertsinidis, A., Zhang, Y., & Chu, S. (2010). Subnanometre single-molecule localization, registration and distance measurements. Nature, 466(7306), 647–651.

8. Roscioli, E., Germanova, T. E., Smith, C. A., Embacher, P. A., Erent, M., Thompson, A. I., et al (2020). Ensemble-level organization of human kinetochores and evidence for distinct tension and attachment sensors. Cell reports, 31(4).

9. Sahl, S. J., Matthias, J., Inamdar, K., Khan, T. A., Weber, M., Becker, S., et al. (2023). Direct optical measurement of intra-molecular distances down to the Angstrom scale. bioRxiv, 2023–07.

10. Suzuki, A., Long, S. K., & Salmon, E. D. (2018). An optimized method for 3D fluorescence co-localization applied to human kinetochore protein architecture. Elife, 7, e32418.

11. Gallego, O., Specht, T., Brach, T., Kumar, A., Gavin, A. C., & Kaksonen, M. (2013). Detection and characterization of protein interactions in vivo by a simple live-cell imaging method. PLoS One, 8(5), e62195.

12. Picco, A., Irastorza-Azcarate, I., Specht, T., Böke, D., Pazos, I., Rivier-Cordey, A. S., et al. (2017). The in vivo architecture of the exocyst provides structural basis for exocytosis. Cell, 168(3), 400–412.

13. Schindelin, J., Arganda-Carreras, I., Frise, E., Kaynig, V., Longair, M., Pietzsch, T., et. (2012). Fiji: an open-source platform for biological-image analysis. Nature methods, 9(7), 676–682.

14. The MathWorks Inc. MATLAB version: 9.13.0 (R2022b), Natick, Massachusetts: The MathWorks Inc. 2002.

15. R Core Team. R: A language and environment for statistical computing. R Foundation for Statistical Computing, 2020, Vienna, Austria.

16. Lu, A. X., Zarin, T., Hsu, I. S., & Moses, A. M. (2019). YeastSpotter: accurate and parameter-free web segmentation for microscopy images of yeast cells. Bioinformatics, 35(21), 4525–4527.

17. Van der Walt, S., Schönberger, J. L., Nunez-Iglesias, J., Boulogne, F., Warner, J. D., Yager, N., … & Yu, T. (2014). scikit-image: image processing in Python. PeerJ, 2, e453.

18. Schroeder, W., Martin, K. M., & Lorensen, W. E. (1998). The visualization toolkit an object-oriented approach to 3D graphics. Prentice-Hall, Inc.

19. Allan, D. B., Caswell, T., Keim, N. C., van der Wel, C. M., & Verweij, R. W. (2021). soft-matter/trackpy: Trackpy v0. 5.0. Zenodo repository.

20. Virtanen, P., Gommers, R., Oliphant, T. E., Haberland, M., Reddy, T., Cournapeau, D., et al. (2020). SciPy 1.0: fundamental algorithms for scientific computing in Python. Nature methods, 17(3), 261–272.

21. Harris, C. R., Millman, K. J., Van Der Walt, S. J., Gommers, R., Virtanen, P., Cournapeau, D., et al. (2020). Array programming with NumPy. Nature, 585(7825), 357–362.

22. McKinney, W. (2010, June). Data structures for statistical computing in python. In Proceedings of the 9th Python in Science Conference (Vol. 445, No. 1, pp. 51–56).

23. Waskom, M., Botvinnik, O., O’Kane, D., Hobson, P., Lukauskas, S., Gemperline, D. C., et al. (2017). mwaskom/seaborn: v0. 8.1. Zenodo.

24. Hunter, J. D. (2007). Matplotlib: A 2D graphics environment. Computing in science & engineering, 9(03), 90–95.

25. Collette, A. (2013). Python and HDF5: unlocking scientific data. “O’Reilly Media, Inc.”.

26. Chollet, F. et al. Keras. GitHub. 2015.

27. Newville, M., Stensitzki, T., Allen, D. B., Rawlik, M., Ingargiola, A., & Nelson, A. (2016). LMFIT: Non-linear least-square minimization and curve-fitting for Python. Astrophysics Source Code Library, ascl-1606.

28. Bradski, G. (2000). The openCV library. Dr. Dobb’s Journal: Software Tools for the Professional Programmer, 25(11), 120–123.

29. Clark, A. Pillow (PIL Fork) Documentation, readthedocs; 2015.

30. Abadi, M. et al. TensorFlow: Large-scale machine learning on heterogeneous systems, software available from tensorflow. org. 2015.

31. Picco, A., & Kaksonen, M. (2017). Precise tracking of the dynamics of multiple proteins in endocytic events. In Methods in cell biology (Vol. 139, pp. 51–68). Academic Press.

