## Supplementary material for "PyF2F: A robust and simplified fluorophore-to-fluorophore distance measurement tool for Protein interactions from Imaging Complexes after Translocation experiments"

Altair C. Hernandez[1], Sebastian Ortiz[1], Laura I. Betancur[1], Radovan Dojčilović[1,2], Andrea Picco[3], Marko Kaksonen[3], Baldo Oliva[1], and Oriol Gallego[1]

[1] Department of Medicine and Life Sciences, Universitat Pompeu Fabra, Barcelona 08005, Catalonia, Spain. [2] Present address: Vinča Institute of Nuclear Sciences, National Institute of the Republic of Serbia, University of Belgrade, P.O. Box 522, 11001 Belgrade, Serbia. [3] Department of Biochemistry, University of Geneva, 1205 Genève, Switzerland

#### Note S1. Bootstrap method for the outlier rejection

Due to the skewed nature of the distance distribution, precise distance estimation requires rejection of possible outliers. We performed the outlier rejection by using the following bootstrap method (for more details, see *Methods* section from Picco A. et al., 2017 (**1**)):

- Each datapoint ($x_i$) is rejected, one at the time, and for each rejection we compute the log-likelihood given an initial estimate of μ and σ:

$$\log L_i(\mu, \sigma) = \sum_{x \in X \setminus \{x_i\}} \log p(x; \mu, \sigma) \qquad \text{(log-likelihood)}$$

  The datapoint that is less likely to belong to the dataset is the one whose rejection gives the worst log-likelihood: $\min_{x_i} \{\log L_i(\mu, \sigma)\}$. This datapoint is rejected and a new estimate of μ and σ is computed.

- The process is iterated by exploring a given percentage of the dataset defined by the user, starting from the largest distances (which are those defining the tail of the distribution, where outliers, if present, are more problematic), has been sampled for rejection. We do not expect to reject as many data points, but ⅓ is a safe parameter to ensure that we had a large sampling of the dataset.

- Two subsequent rejections $i$ and $i+1$ will give two estimations of μ. Their difference, $\delta\mu_i = \mu_{i+1} - \mu_i$, will decrease when most outliers are rejected and the score will thus be maximal:

$$p_{\delta\mu_i} = \frac{1/\delta\mu_i}{\sum_j 1/\delta\mu_j},\ p_{\delta\sigma_i} = \frac{1/\delta\sigma_i}{\sum_j 1/\delta\sigma_j} \qquad \text{(scores for each iteration)}$$

The μ, σ and the ensemble of data points that are retained after all these iterations are those that maximise a scoring function defined as:

$$S(p_{\delta\mu_i}, p_{\delta\sigma_i}) =- p_{\delta\mu_i}\log\left(p_{\delta\mu_i}\right) - p_{\delta\sigma_i}\log\left(p_{\delta\sigma_i}\right) \qquad \text{(scoring function)}$$

The total score $S(p_{\delta\mu_i}, p_{\delta\sigma_i})$ will be maximal when both scores $p_{\delta\mu_i}$ and $p_{\delta\sigma_i}$ will be similarly maximised ( $\max_i (S(p_{\delta\mu_i}, p_{\delta\sigma_i}))$).

#### Table S1. Comparison of distance estimations

PyF2F performance was evaluated by comparing the distances estimated from a published PICT dataset (**1**) and comparing the results obtained with PyF2F (**μ**_PyF2F - **SE$_\mu$**_PyF2F) with the published ones (**μ**_Picco - **SE$_\mu$**_Picco). We compared 45 distance estimations corresponding to the separation between the anchor labelled with RFP and C-terminally-tagged exocyst subunits (GFP_C); 33 distance estimations corresponding to the separation between the anchor labelled with RFP and N-terminally-tagged exocyst subunits (GFP_N); and 6 inter-assembly distance estimations corresponding to the separation between the anchor labelled with RFP and Sec2 fused to GFP (Sec2_GFP_C).

**μ:** Distance estimation.
**SE$_\mu$:** Standard error of the distance estimation.

| Bait | Prey | μ_Picco (nm) | SE$_\mu$_Picco (nm) | μ_PyF2F (nm) | SE$_\mu$_PyF2F (nm) |
|---|---|---|---|---|---|
| Sec3-FRB | Sec5_GFP_C | 19.92 | 1.37 | 23.03 | 1.21 |
| Sec3-FRB | Sec6_GFP_C | 20 | 2.73 | 20.82 | 2.64 |

| | | | | | |
|---|---|---|---|---|---|
| Sec3-FRB | Sec8_GFP_C | 18.72 | 2.4 | 23.98 | 1.97 |
| Sec3-FRB | Sec10_GFP_C | 24.52 | 2.47 | 25.09 | 3.84 |
| Sec3-FRB | Sec15_GFP_C | 21.64 | 3.28 | 18.8 | 2.95 |
| Sec3-FRB | Exo70_GFP_C | 16.41 | 2.79 | 17.8 | 1.34 |
| Sec3-FRB | Exo84_GFP_C | 19.79 | 3.07 | 21.89 | 4.54 |
| Sec6-FRB | Sec3_GFP_C | 26.96 | 2.48 | 23.62 | 2.31 |
| Sec6-FRB | Sec5_GFP_C | 24.49 | 2.76 | 30.02 | 2.96 |
| Sec6-FRB | Sec15_GFP_C | 20.55 | 4.88 | 25.51 | 3.83 |
| Sec8-FRB | Exo70_GFP_C | 22.04 | 2.02 | 20.98 | 2.8 |
| Sec8-FRB | Exo84_GFP_C | 17.61 | 2.35 | 18.59 | 3.15 |
| Sec6-FRB | Exo70_GFP_C | 22.71 | 2.49 | 20.57 | 2.9 |
| Sec6-FRB | Exo84_GFP_C | 24.67 | 3.02 | 28.75 | 3.12 |
| Sec8-FRB | Sec3_GFP_C | 20.9 | 2.36 | 21.15 | 5.46 |
| Sec8-FRB | Sec5_GFP_C | 20.15 | 1.98 | 22.72 | 1.97 |
| Sec8-FRB | Sec6_GFP_C | 22.87 | 2.24 | 24.82 | 2.69 |
| Sec8-FRB | Sec10_GFP_C | 20.68 | 1.73 | 21.97 | 2.62 |
| Sec15-FRB | Sec3_GFP_C | 21.99 | 2.86 | 23.06 | 3.42 |
| Sec15-FRB | Sec5_GFP_C | 21.84 | 2.01 | 19.68 | 1.57 |
| Sec10-FRB | Sec3_GFP_C | 22.5 | 2.79 | 24.86 | 4.79 |
| Sec10-FRB | Sec5_GFP_C | 25.85 | 2.51 | 25.43 | 2.78 |
| Sec10-FRB | Sec8_GFP_C | 20.17 | 1.71 | 24.73 | 2.77 |
| Sec10-FRB | Sec15_GFP_C | 23.25 | 1.88 | 20.79 | 2.56 |
| Sec10-FRB | Exo70_GFP_C | 24.29 | 2.43 | 27.93 | 2.63 |
| Sec10-FRB | Exo84_GFP_C | 25.14 | 2.41 | 26.67 | 3.26 |
| Sec15-FRB | Sec6_GFP_C | 21.75 | 2.73 | 25.24 | 2.04 |
| Exo70-FRB | Sec6_GFP_C | 25.87 | 1.98 | 22.69 | 2.3 |
| Exo70-FRB | Sec8_GFP_C | 29.3 | 1.74 | 34.96 | 1.58 |
| Exo70-FRB | Sec10_GFP_C | 31.69 | 1.94 | 32.66 | 2.08 |
| Sec15-FRB | Sec8_GFP_C | 17.72 | 2.24 | 17.21 | 1.94 |
| Sec15-FRB | Sec10_GFP_C | 13.91 | 3.35 | 21.37 | 2.35 |
| Sec15-FRB | Exo70_GFP_C | 22.12 | 1.72 | 25.91 | 2 |

| | | | | | |
|---|---|---|---|---|---|
| Sec15-FRB | Exo84_GFP_C | 19.68 | 2.48 | 16.42 | 1.82 |
| Exo70-FRB | Sec3_GFP_C | 24.02 | 1.3 | 25.35 | 1.45 |
| Exo70-FRB | Sec5_GFP_C | 26.88 | 1.36 | 24.12 | 1.77 |
| Exo70-FRB | Sec15_GFP_C | 32.73 | 1.73 | 32.92 | 3.14 |
| Exo84-FRB | Sec15_GFP_C | 20.49 | 2.77 | 24.59 | 3.1 |
| Exo84-FRB | Exo70_GFP_C | 24.78 | 1.76 | 25.48 | 2.22 |
| Exo70-FRB | Exo84_GFP_C | 29.09 | 1.77 | 34.25 | 1.38 |
| Exo84-FRB | Sec3_GFP_C | 21.04 | 1.72 | 21.47 | 3.07 |
| Exo84-FRB | Sec5_GFP_C | 19.5 | 2.31 | 19.73 | 2.71 |
| Exo84-FRB | Sec6_GFP_C | 24.79 | 2.57 | 29.45 | 3.04 |
| Exo84-FRB | Sec8_GFP_C | 20.6 | 1.53 | 21.19 | 2.49 |
| Exo84-FRB | Sec10_GFP_C | 18.6 | 2.36 | 25.91 | 3.31 |
| Exo70-FRB | Sec5_GFP_N | 20.46 | 1.67 | 22.42 | 2.27 |
| Sec3-FRB | Sec5_GFP_N | 22.51 | 1.41 | 19.73 | 1.79 |
| Sec3-FRB | Sec6_GFP_N | 20.56 | 2.06 | 20.72 | 1.86 |
| Sec3-FRB | Sec10_GFP_N | 20.56 | 3.65 | 22.95 | 3.03 |
| Sec3-FRB | Sec15_GFP_N | 23.65 | 3.61 | 25.64 | 6.05 |
| Sec6-FRB | Sec3_GFP_N | 21.09 | 4.19 | 23.31 | 2.63 |
| Sec8-FRB | Sec3_GFP_N | 24 | 2.02 | 20.66 | 1.82 |
| Sec8-FRB | Sec5_GFP_N | 22.47 | 2.26 | 20.91 | 0.83 |
| Sec8-FRB | Sec6_GFP_N | 18.92 | 1.44 | 22.94 | 1.83 |
| Sec8-FRB | Sec10_GFP_N | 27.37 | 2.51 | 28.02 | 2.05 |
| Sec15-FRB | Sec3_GFP_N | 23.91 | 2.83 | 24.06 | 2.45 |
| Sec15-FRB | Sec5_GFP_N | 23.83 | 3.63 | 30.03 | 2.97 |
| Sec10-FRB | Sec5_GFP_N | 15.55 | 4.82 | 27.95 | 3.73 |
| Sec10-FRB | Sec6_GFP_N | 23.48 | 2.72 | 23.72 | 0.32 |
| Sec10-FRB | Sec15_GFP_N | 16.78 | 2.21 | 17.98 | 2.97 |
| Sec15-FRB | Sec6_GFP_N | 27.51 | 2.63 | 26.5 | 2.93 |
| Exo70-FRB | Sec6_GFP_N | 25.29 | 1.81 | 24.39 | 1.5 |
| Sec15-FRB | Sec10_GFP_N | 21.47 | 2.83 | 22.92 | 2.7 |
| Exo70-FRB | Sec3_GFP_N | 19.72 | 2.36 | 19.35 | 1.81 |

| | | | | | |
|---|---|---|---|---|---|
| Exo70-FRB | Sec15_GFP_N | 21.05 | 1.8 | 25.13 | 2.09 |
| Exo70-FRB | Exo84_GFP_N | 18.38 | 1.63 | 17.16 | 2.38 |
| Exo84-FRB | Sec3_GFP_N | 19.45 | 2.11 | 21.86 | 2.62 |
| Exo84-FRB | Sec5_GFP_N | 23.92 | 1.95 | 28.39 | 2.49 |
| Exo84-FRB | Sec6_GFP_N | 27.15 | 1.77 | 25.26 | 1.12 |
| Exo84-FRB | Sec10_GFP_N | 23.47 | 2.74 | 30.64 | 0.78 |
| Sec3-FRB | Exo70_GFP_N | 23.28 | 2.32 | 25.82 | 2.13 |
| Sec8-FRB | Exo84_GFP_N | 23.34 | 1.74 | 23.89 | 1.78 |
| Sec10-FRB | Exo84_GFP_N | 15.36 | 2.37 | 31.89 | 3.3 |
| Sec8-FRB | Exo70_GFP_N | 23.26 | 1.8 | 21.66 | 3.02 |
| Sec10-FRB | Exo70_GFP_N | 22.04 | 3.21 | 34.08 | 3.83 |
| Sec3-FRB | Exo84_GFP_N | 22.55 | 2.01 | 21.58 | 2.48 |
| Exo84-FRB | Exo70_GFP_N | 18.92 | 3.25 | 24.71 | 3.36 |
| Sec15-FRB | Exo84_GFP_N | 24.32 | 2.23 | 22.71 | 2.19 |
| Sec3-FRB | Sec2_GFP_C | 65.22 | 5.13 | 78.73 | 5.65 |
| Sec6-FRB | Sec2_GFP_C | 68.31 | 9.69 | 68 | 5.1 |
| Sec10-FRB | Sec2_GFP_C | 66.69 | 6.51 | 68.96 | 7.05 |
| Sec15-FRB | Sec2_GFP_C | 69.06 | 15.41 | 59.76 | 5.33 |
| Exo70-FRB | Sec2_GFP_C | 78.8 | 5.18 | 81.57 | 5.03 |
| Exo84-FRB | Sec2_GFP_C | 64.87 | 10.03 | 55 | 13.07 |

### REFERENCES

1. Picco, A., & Kaksonen, M. (2017). Precise tracking of the dynamics of multiple proteins in endocytic events. In *Methods in cell biology* (Vol. 139, pp. 51-68). Academic Press.
